# Enhancing tandem MS sensitivity and peptide identification via ion pre-accumulation in an Orbitrap mass spectrometer

**DOI:** 10.1101/2025.02.20.639277

**Authors:** Florian Harking, Ulises H. Guzman, Julia Kraegenbring, Hamish Stewart, Konstantin Aizikov, Heiner Koch, Kyle L. Fort, Alexander Harder, Jesper V. Olsen

## Abstract

High-throughput mass spectrometry-based proteomics has gained increasing interest for both academic and industrial applications. As implementation of faster gradients has facilitated higher sample throughput, mass spectrometers must adapt to shorter analysis times by enhancing scanning speed and sensitivity. For Orbitrap™ mass spectrometers, faster scan rates are constrained by the need for sufficient ion accumulation time, particularly given limitations on duty cycle at high repetition rate, and transient length, which determines analyzer sensitivity and resolving power. In this context, implementing alternative ion scheduling and better ion signal-processing strategies are needed to unleash the speed of these instruments. Here, we introduce a new scanning strategy termed pre-accumulation, which enables the storage of ions in the bent flatapole in parallel to the operation of the C-trap/IRM, leading to a significant improvement in ion beam utilization and enabling for the first time scanning speeds of >70 Hz on hybrid Orbitrap instruments. The combination of pre-accumulation and increased scan speeds notably enhances peptide and protein group identifications for short LC gradients and improves sensitivity for high-throughput applications. These benefits were further amplified when coupled with the full mass range phase-constrained spectrum deconvolution method (ΦSDM), especially for fast, lower-resolving Orbitrap measurements used with short LC gradients. Overall, we demonstrate that pre-accumulation of ions in the bent flatapole offers distinct advantages, particularly for conditions with reduced signal input. Since no hardware changes are required, this approach is highly attractive for Orbitrap mass spectrometers operated with fast MS/MS acquisition methods.

## Introduction

Mass spectrometry (MS)-based proteomics has become the method of choice for comprehensive and unbiased quantification and identification of thousands of proteins across a variety of biological samples^1^. Among the mass spectrometers available, hybrid Orbitrap™ instruments such as the Thermo Scientific Orbitrap Exploris™ 480 Mass Spectrometer, have become workhorses in modern day proteomics applications^13^. However, exploring complex biological systems requires scaling the analysis of these samples, which often involves implementing short chromatographic gradients in liquid chromatography tandem mass spectrometry (LC-MS/MS) workflows^2,3^ and automated sample preparation protocols^4,5^ to enhance the throughput of proteomics experiments. As a result, it is essential for MS instruments to possess the capability of measuring eluting peptides at high scanning speeds^6^. Technological advancements in both hardware and software have been pivotal in facilitating faster acquisition methods, thereby improving the throughput of proteomic experiments^7–11^. In recent years, the proteomics community has seen the development of mass spectrometers with scanning speeds ranging from 100 to 300 Hz^12^. However, while hybrid Orbitrap instruments provide several advantages such as their compact design, exceptional resolving power, sensitivity and robustness making them widely utilized in proteomics laboratories worldwide, they are limited to scanning speeds of < 50 Hz. This is in part due to the fact that higher scan speeds constrain the Orbitrap transient length duration, limiting the resolving power and sensitivity of the analyzer^14,15^. Moreover, maintaining the sensitivity of MS methods depends on having a sufficient number of ions in the instrument, which is proportional to the injection time^16^. This aspect is crucial, as the shorter injection times required for full acquisition parallelization of the instrument when operated at fast scanning speeds (>50 Hz) can affect the quality of the acquired fragment ion spectra^17,18^. Importantly, this limitation is not due to the Orbitrap analyzer itself, but due to duty cycle losses caused by fixed timing overheads associated with the operation of the C-Trap and Ion Routing Multipole (IRM) that accumulate, prepare and inject ions into the Orbitrap analyzer. Balancing the Orbitrap transient length and injection time allows the instrument to operate in full parallelization mode, where injection time may approach the length of the transient^19^, excepting overheads. To address these challenges, we evaluated a feature integrated into a modified Orbitrap Exploris 480 instrument control software that enables the trapping and accumulation of ions in the bent flatapole, upstream of the C-Trap/IRM, in parallel with transient acquisition. Additionally, we employed the phase-constrained spectrum deconvolution method (ΦSDM)^20^ for transient analysis at full MS and MS/MS range, which allowed us to achieve more than twofold higher mass resolving power compared to conventional processing with the enhanced Fourier Transform^21^ (eFT) at equivalent transient lengths^22,23^. The integration of these two features facilitated the use of even shorter transient lengths, enabling MS/MS acquisition speeds of up to 70 Hz and the generation of high-quality fragmented spectra, allowing more rapid scanning of the chromatographic peaks, ultimately enhancing the overall performance of the instrument.

## Results

### Pre-accumulation feature: Proof of Concept

As mentioned above, despite advancements in hybrid Orbitrap instrumentation, the maximum acquisition rate is constrained to ∼44 Hz (7.5k Resolution at m/z 200). This limitation arises from time overheads associated with the operation of the C-trap and IRM. Briefly, in a typical mass spectrometry experiment, ions are injected into the C-trap, then transferred to the IRM for cooling, and subsequently transferred back to the C-trap for orthogonal ejection to the Orbitrap analyzer. During this process, no additional ions can be accumulated, leading to a loss of the ion beam at the charge detector (**Fig. 1a**). We hypothesized that optimizing the instrument’s operation by circumventing these time overheads could enhance overall instrument performance. Therefore, we leveraged the bent flatapole, which serves as a curved ion transfer guide between the ion source and the C-Trap/IRM. This device can be repurposed as an ion trap due to its quadrupole structure and exit lens aperture, which features a superimposed direct current (DC) gradient independent of the quadrupole mass filter. By adjusting the DC voltage applied to the exit lens, we could effectively control the trapping and releasing of ions by switching from a trapping potential (+10V) to a transmitting potential (−35V). This enabled a suitable method (Pre-accumulation) that bypassed the C-trap/IRM reset dead time, providing up to 10 ms of additional ion accumulation time available for each HCD-MS/MS scan, when operating the system at scanning speed of 44 Hz with an Orbitrap resolution of 7,500 at m/z 200 (**Fig. 1b)**. To test the feasibility of this method, we acquired MS/MS spectra of the isolated MRFA peptide using a maximum injection time of 10 ms Orbitrap transient with the Pre-accumulation feature on and off (**Fig. 1c**). The results showed a 125% increase in the overall MS/MS signal with the pre-accumulation feature activated, while preserving the relative intensities of the fragments. This enhancement could be attributed to the doubling of effective injection time when the Pre-accumulation feature was enabled, resulting in greater number of ions available for detection.

**Fig. 1.**
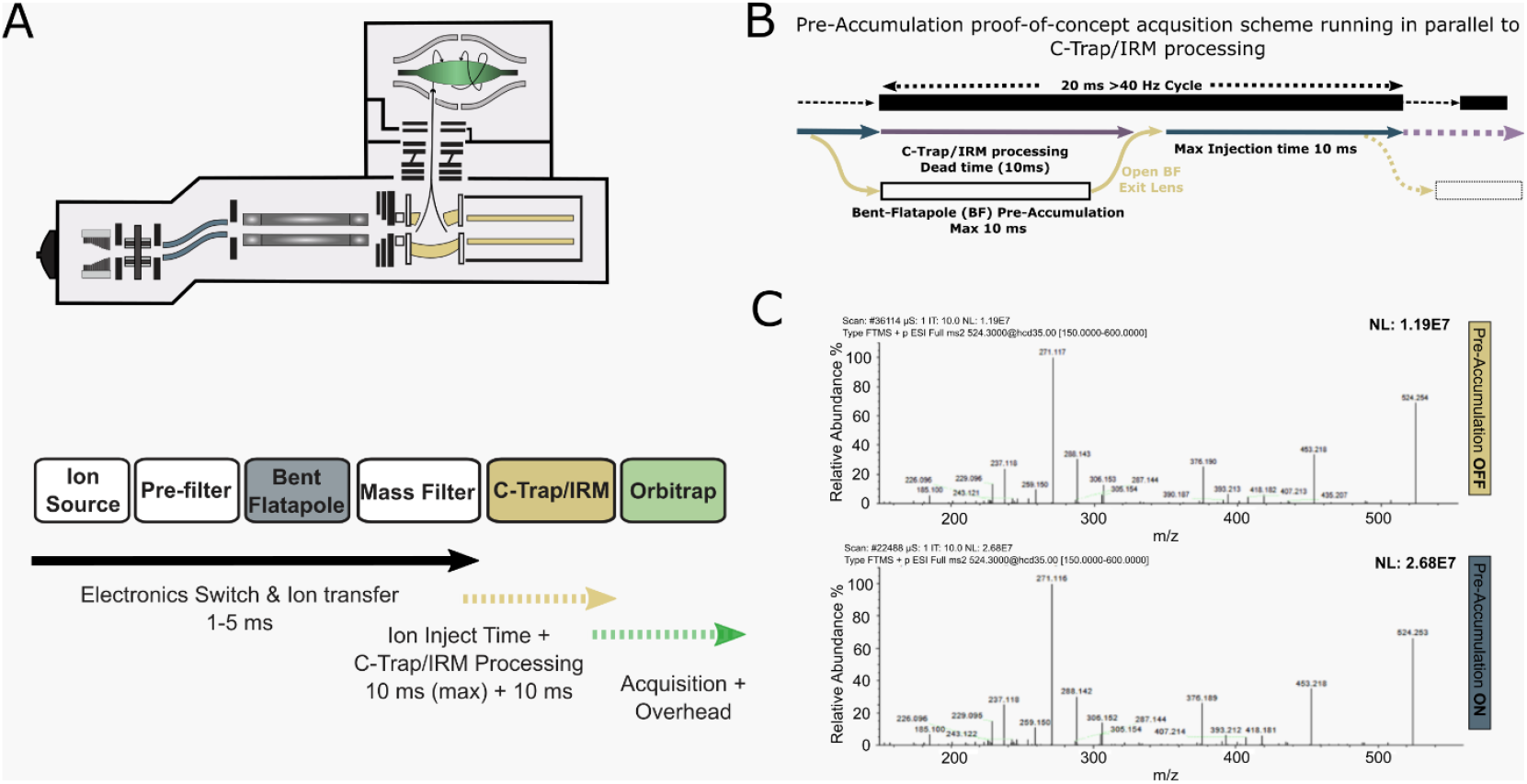
Proof-of-concept of the ion pre-accumulation feature. **A)** Orbitrap Exploris™ 480 mass spectrometer layout (top). Operation scheme of the Orbitrap Exploris™ 480 mass spectrometer, depicting the ion pre-accumulation process within the bent flatapole while the C-trap/IRM is engaged (bottom). **B)** Analysis of the pre-Accumulation acquisition scheme in parallel to the C-Trap/IRM Processing. The pre-accumulation feature enables to accumulate the ion bean at the bent flatapole until the C-trap resets - enabling additional up to 10 ms fill time at a resolution of 7500 resolution @ m/z 200 (40 Hz scanning speed). **C)** MS2 spectra of the MRFA peptide (524.3 m/z) collected at maximum injection time of 10 ms, normalized collision energy (NCE) 35 and 7500 resolution with the pre-accumulation feature off (upper) and on (bottom).

### Effect of Pre-accumulation feature on rapid MS2 scanning on Data-dependent acquisition mode

The scanning speed of mass spectrometers is a vital parameter in proteomics research. Enhanced scanning speeds enable rapid scan cycles, which allows for the detection of a broader dynamic range of ion signals^19^ and results in more scans across a chromatographic peak, which facilitates high-throughput applications^24^. However, achieving rapid MS/MS scanning speed requires shorter transients and reduced fill times, leading to a proportional decrease in ion flux. This reduction adversely impacts scan quality, as fewer ions are measured by the detector^16,25^. A common solution to this issue is to analyze high-input loads. Therefore, to assess the effect of the additional injection time delivered by the Pre-accumulation feature, we analyzed a dilution series of a HeLa tryptic digest in data-dependent acquisition mode (DDA) with short 8-min online LC gradients (180 SPD). In addition, to take full advantage of the Pre-accumulation feature, we employed rapid scan speeds of up to 70 Hz by constraining the Orbitrap transient length to 8 ms. Reassuringly, we observed that increasing the scanning speed from 22 Hz to 70 Hz resulted in a proportional increase in the total number of MS/MS scans per raw file (**Fig. 2a**). Our results showed an increase in both peptide and protein group identifications when the Pre-accumulation feature was enabled, independent of the input amount up to 44 Hz scanning speed. Furthermore, to mitigate the reduction in observed resolution due to shorter transient lengths, we employed ΦSDM^20^ for processing at full MS and MS/MS spectra across the entire mass ranges. Notably, the most significant benefits were observed when the Pre-accumulation feature was combined with ΦSDM (ΦSDM+Preacc), particularly at high scanning speeds (**Fig. 2b)**. Notably, our results showed a ∼20% and ∼40 % increase in protein group and peptide identifications quantified in at least 70% of the data set, when ΦSDM+Preacc at 70 Hz is compared to the fastest standard scanning condition (44 Hz) with 500 ng input. Moreover, when comparing ΦSDM+Preacc at 70 Hz with the slowest standard scanning condition (22 Hz), we observed increases of approximately 130% and 100 % in peptide and protein group identifications, respectively, also with 500 ng input under the same quantification criteria.

**Fig. 2.**
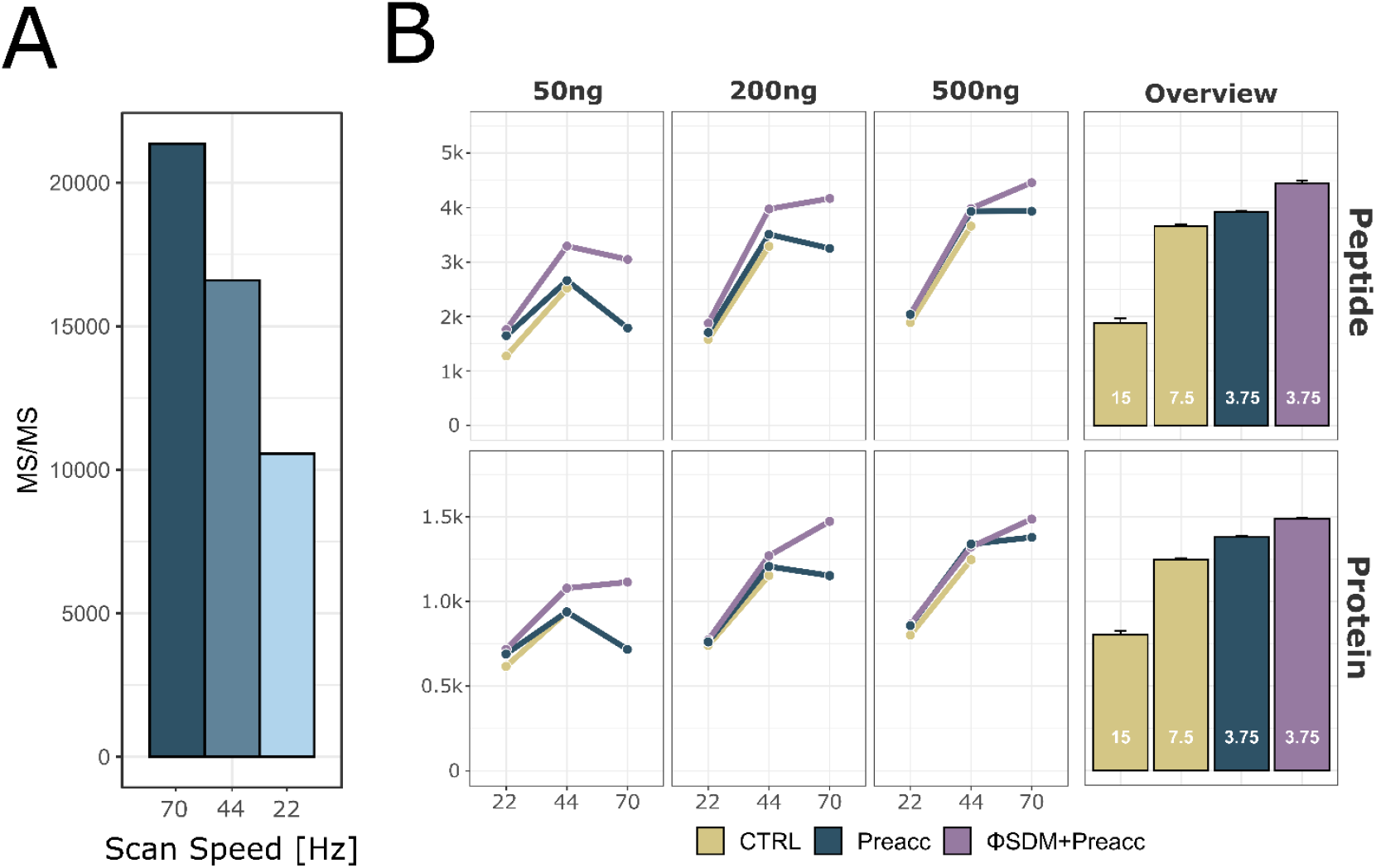
Pre-accumulation feature increases the identification rate of rapid MS2 scanning in Data-dependent acquisition mode. **A)** MS2 scan number across different scanning speed for 8-min chromatographic gradient in Data-dependent acquisition mode (500ng input). **B)** Overview of peptide (top) and protein (bottom) identifications at different scans speed (22, 44 and 70Hz) across different Hela tryptic peptide inputs and three operating modes (CTRL: default operation mode, Preacc: Pre-accumulation feature enabled and ΦSDM+Preacc: Pre-accumulation and ΦSDM (MS1-MS2) features enabled, Best: input 500ng). Chromatographic gradient was constant (8-min, 180 SPD). Data presented for Peptides/Protein groups quantified at minimum in 70% of the data set.

### Improving the sensitivity of Data-independent acquisition mode

Data-independent acquisition (DIA) has emerged as the method of choice for quantitative high-throughput analysis of samples, primarily due to its exceptional sensitivity and its combination with short single-shot LC gradients^8^. Given that both identification and quantification rely on the fragmentation spectra^26,27^, we hypothesized that the improved signal intensity in MS/MS spectra observed for the MRFA peptide when the Pre-accumulation feature was enabled, could benefit acquisitions in DIA mode and short LC gradients (200 SPD; 5.6 min active gradients). Therefore, we analyzed a dilution series of HeLa tryptic peptides spanning from 5 to 500 ng input amounts. We observed an increase in protein group and peptide identifications when the Pre-accumulation feature was enabled, regardless of the input amount. This effect was particularly notable at low input amounts (<50 ng) compared to the control scanning condition. Conversely, enabling only the full mass range ΦSDM at MS2 (ΦSDM-MS2) resulted in increased protein groups and peptide identifications across the entire dilution series, with better performance at high input amounts compared to the Pre-accumulation and control scanning conditions (**Fig. 3a**). Notably, when both the Pre-accumulation and ΦSDM-MS2 features were enabled, we observed increases in peptide and protein group identifications compared to the control, ΦSDM-MS2 and Pre-accumulation only scanning methods for input amounts under 50ng. Furthermore, under optimal conditions (25ng tryptic peptide input), there was an increase of over 30% in the identification of peptide and protein groups (**Fig. 3b**).The increase in identifications observed under both conditions (Pre-accumulation and ΦSDM-MS2+Preacc) could be attributed to the overall enhancement of the MS/MS signal relative to the standard scanning condition (**Fig. 3c**), consistent with our previous findings.

**Fig. 3.**
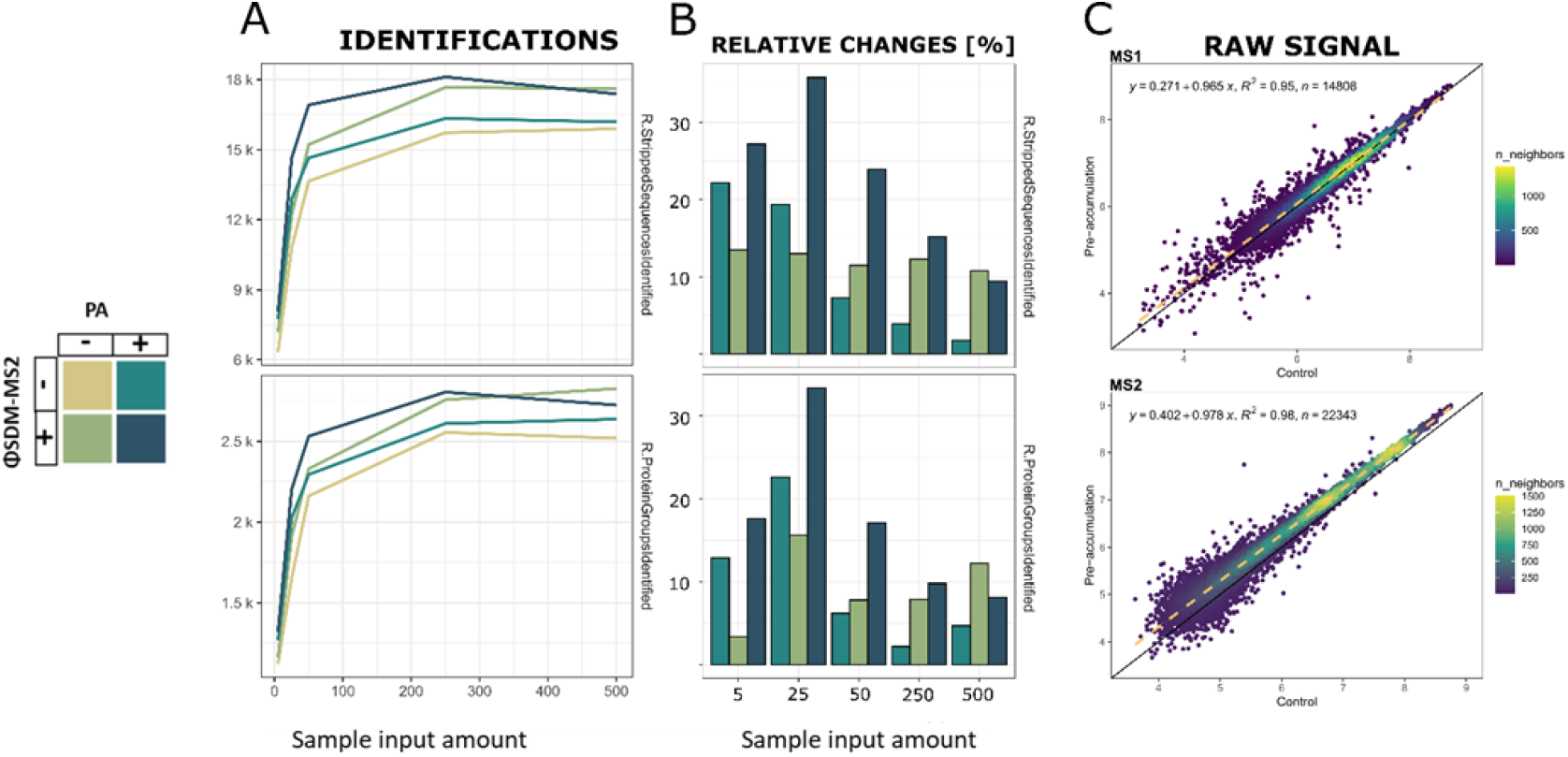
Pre-accumulation feature increases sensitivity and performance of Data-independent acquisition scanning schemes. **A)** Identification numbers at peptide (top) and protein group level (bottom) according to different input amounts and operating modes. (PA : Pre-accumulation, ΦSDM-MS2, +; on and -; off). **B)** Relative identifications at peptide level (top) and protein group level (bottom) compared to respective control (PA; off, ΦSDM-MS2; off) across different input amounts. **C)** Comparison of raw ion signal at the MS1 level (top) and MS2 level (bottom) between control and pre-accumulation conditions.

## Discussion

In this technical note, we present the implementation of an enhanced MS/MS Pre-accumulation method for Orbitrap instruments, which can be employed routinely in proteomics experiments. This approach facilitates the ion trapping and accumulation to occur concurrently with the C-trap/IRM operation in an Orbitrap Exploris 480 MS. By accumulating ions in this region when in typical operation they would be lost, we enhance ion utilization and increase the ion flux reaching the MS detector, as shown by the MS2 intensity improvements for both infused FlexMix™ and LC-MS/MS experiments of HeLa tryptic digests when Pre-accumulation was enabled. This enhanced number of ions available for MS/MS acquisition is particularly interesting in combination with rapid scanning speed methods and short gradients since such scanning methods require the right balance between Orbitrap resolution and injection time, limiting the Orbitrap scan rate to < 50 Hz. As the Pre-accumulation feature allows for an extended accumulation time of ions for MS2 spectra while maintaining the full parallelization of the instrument, it is feasible to increase the Orbitrap scanning rate to >70 Hz by reducing the MS2 resolution to 3,750 at 200 m/z. Although, the 44 Hz DDA scanning method with the Pre-accumulation feature enabled demonstrated an increment in the identification rate compared to the control scanning condition, this improvement becomes less evident as the input amount decreases. Notably, this effect is more pronounced when the instrument was operated at >70 Hz, which was only possible to run with the Pre-accumulation feature switched on. This trend is expected, as the use of short Orbitrap transient lengths compromises the sensitivity and resolution achieved by the detector, since these factors scale with the Orbitrap transient length^28,29^. Consequently, this most likely reduces the number of precursor candidates observed at the MS1 level that are necessary for fragmentation in DDA methods. To address this, we employed the full mass range ΦSDM feature at MS1 and MS2 using short transient length methods, which enables to obtain higher mass resolution at the same transient length when compared to eFTs^20,22^.

Enabling the ΦSDM and the Pre-accumulation features together, we boost the identification rate of 44 Hz DDA methods for inputs under 500 ng. Conversely, the ΦSDM+Preacc method at 70 Hz exhibited enhanced performance over other methods for input amounts exceeding 50 ng. This phenomenon can be attributed to the fact that high input amounts can compensate for the reduced ion flux reaching the detector. This reduction results from the short injection times required for parallelizing fast scanning DDA methods that employ low MS1 resolutions, generating fragment spectra with enough quality for confident peptide identification^16,18,24^. However, the improvement in spectrum quality provided by extended injection times from the Pre-accumulation feature alone is insufficient to enhance DDA methods. This is because, in these methods, MS1 and MS2 are interconnected; thus, enabling the ΦSDM feature across full mass range is essential for resolving MS1 and MS2 peaks at short transient lengths. This capability generates more candidates for fragmentation and enhances the resolution of fragment peaks, improving MS/MS spectra quality. In contrast, DIA methods directly benefit from the Pre-accumulation feature, as they rely on MS/MS spectra for identification and quantification. Notably, the advantages of Pre-accumulation are more pronounced at low input amounts, and this effect is further amplified when Pre-accumulation is combined with ΦSDM-MS2. This synergic effect can be attributed to the extended ion injection coming from the Pre-accumulation feature and the improved resolution and more accurate mass-to-charge ratios provided by the ΦSDM feature^20^. Additionally, it is important to note that the contributions of both features (Pre-accumulation and ΦSDM-MS2) vary across the tested input amount range, influencing the identification rate differently.

The Pre-accumulation feature has a greater impact at low loads, while ΦSDM-MS2 enhances the identification rate across all conditions. This demonstrates the complementarity of the two technologies: low input amounts benefit the most from extended injections times, as more ions are allowed to reach the detector, whereas this benefit diminishes as higher input amounts. In contrast, ΦSDM-MS2 improves the spectra quality, especially at short transient lengths, enhancing the data across all tested input amounts. Overall, we demonstrated the feasibility of implementing a Pre-accumulation feature repurposing the bent flatapole as an ion-trap to leverage the C-trap/IRM reset dead time. This significantly enhanced the performance of hybrid Orbitrap instruments and enables MS/MS acquisition methods at >70 Hz. This method is highly attractive to implement in high-throughput proteomics applications, where fast scanning speeds are needed.

## Materials and Methods

### Sample preparation

HeLa S3 cervical carcinoma cells were cultured under conditions recommended by the manufacturer and harvested at 70% confluence, as determined by visual inspection. Cells were rinsed twice with PBS buffer and subsequently lysed using boiling 1% SDS buffer. The lysates were transferred to Eppendorf vials and stored at -80°C until further use. Prior to processing, samples were thawed at 50°C and subjected to sonication using a stick sonication device. The lysate was then centrifuged at 1,000 x g for 1 minute, and the supernatant was collected for further analysis. HeLa lysates were digested using protein aggregation capture (PAC) on a Kingfisher robot^30,31^. The resulting peptide digests were acidified with formic acid to a final concentration of 1%. Solid phase extraction (SPE) was performed using SepPak 50 mg C18 cartridges on a vacuum manifold. Finally, the peptide mixture was quantified using a Nanodrop at 280 nm before being concentrated via SpeedVac centrifugation. The dried peptides were stored at -20°C until further analysis.

### Mass spectrometry analysis

The Orbitrap Exploris™ 480 mass spectrometer was operated using prototype software from Thermo Fisher Scientific, which enabled the trapping of ions in the Bent Flatapole for fragment scans (MS2). When full mass range ΦSDM processing was required, the spectra were analyzed on a dedicated external computer equipped with GPU cards. The number of iterations was set to 150, with a noise threshold for peak detection established at 1.4. During DDA experiments ΦSDM at full MS and MS/MS was used (ΦSDM) while DIA experiments ΦSDM was used in MS/MS only (ΦSDM-MS2). Phases were pre-calibrated across the entire mass range using the FlexMix™ solution. DDA runs were conducted using the Vanquish Neo LC system in conjunction with the Orbitrap Exploris™ 480 Mass Spectrometer. Briefly, an 8-minute gradient was implemented at a flow rate of 750 nl/min. The gradient increased from 4% to 22.5% B from 0 to 3.7 minutes, then to 45% B by 5.5 minutes, and finally escalated to 99% B for the remainder of the gradient. The DDA method operated at a resolution of 45,000 for MS1, covering a mass range of 375-1200 m/z, utilizing a Top40 approach with a minimum intensity threshold of 5000. MS2 resolution varied among 3,750, 7,500, and 15,000. DIA runs were performed on an EvoSep One LC system coupled to an Orbitrap Exploris™ 480 mass spectrometer using the 200 SPD (5.6 min gradient). Briefly, the method employed MS1 resolution of 120,000 and a maximum injection time of 5 ms while MS2 scans were conducted at a resolution of 15,000, with a maximum injection time of 22 ms, covering a mass range of 361 to 1033 m/z and utilizing a window size of 13.7, resulting in 49 windows. Finally, the samples were prepared using specially designed tips, with load conditions of 0, 5, 25, 50, 250, and 500 ng of Hela tryptic peptides, each tested in triplicate. The experimental conditions were randomized, progressing from lower to higher loads.

### Data analysis

DDA data was analyzed using MaxQuant version 3.6.1^32^, employing a fully reviewed SwissProt library containing 20,223 entries. Each RAW file was annotated individually. For DIA data, Spectronaut version 18. was utilized, also with a fully reviewed SwissProt library of the same size. All files were processed collectively in directDIA mode, using the “Method evaluation” setting, with each condition annotated separately and cross-run normalization turned off. Subsequent data processing and analysis were performed in R version 4.3.1, utilizing the data.table, tidyverse^33^, and ggpubr packages. Data visualization was conducted using the ggplot2 package^34^. Raw data extraction from Thermo RAW files was achieved in R through the rawrr package^35^.

## Acknowledgments

We would like to acknowledge Prof. Dr. Alexander Makarov, Dr. Dmitry Grinfeld, Oliver Lange, Dr. Arne Kreutzmann, and Dr. Daniel Mourad from Thermo Fisher Scientific for fruitful discussions that contributed to the success of this manuscript.

## Data and materials availability

Mass spectrometry data are deposited in the ProteomeXchange^13^ Consortium: XXXXXX

## References

1. Cui, M., Cheng, C. & Zhang, L. High-throughput proteomics: a methodological minireview. Lab. Investig. 102, 1170–1181 (2022).

2. Bache, N. et al. A novel LC system embeds analytes in pre-formed gradients for rapid, ultra-robust proteomics. Mol. Cell. Proteomics 17, 2284–2296 (2018).

3. Kuster, B. et al. Robust microflow LC-MS/MS for proteome analysis: 38 000 runs and counting. Anal. Chem. 93, 3686–3690 (2021).

4. Kverneland, A. H. et al. Fully Automated Workflow for Integrated Sample Digestion and Evotip Loading Enabling High-Throughput Clinical Proteomics. Mol. Cell. Proteomics 23, 100790 (2024).

5. Liang, Y. et al. Fully automated sample processing and analysis workflow for low-input proteome profiling. Anal. Chem. 93, 1658–1666 (2021).

6. Proteomics - 2022 - Messner - Mass spectrometry-based high-throughput proteomics and its role in biomedical studies and.pdf.

7. Messner, C. B. et al. Ultra-fast proteomics with Scanning SWATH. Nat. Biotechnol. 39, 846–854 (2021).

8. Bekker-Jensen, D. B. et al. A compact quadrupole-orbitrap mass spectrometer with FAIMS interface improves proteome coverage in short LC gradients. Mol. Cell. Proteomics 19, 716–729 (2020).

9. Meier, F., Geyer, P. E., Virreira Winter, S., Cox, J. & Mann, M. BoxCar acquisition method enables single-shot proteomics at a depth of 10,000 proteins in 100 minutes. Nat. Methods 15, 440–448 (2018).

10. Demichev, V., Messner, C. B., Vernardis, S. I., Lilley, K. S. & Ralser, M. DIA-NN: neural networks and interference correction enable deep proteome coverage in high throughput. Nat. Methods 17, 41–44 (2020).

11. Wang, Z. et al. High-throughput proteomics of nanogram-scale samples with Zeno SWATH MS. Elife 11, (2022).

12. Peters-Clarke, T. M., Coon, J. J. & Riley, N. M. Instrumentation at the Leading Edge of Proteomics. Anal. Chem. 96, 7976–8010 (2024).

13. Vizcaíno, J. A. et al. ProteomeXchange provides globally coordinated proteomics data submission and dissemination. Nat. Biotechnol. 32, 223–226 (2014).

14. Zubarev, R. A. & Makarov, A. Orbitrap mass spectrometry. Anal. Chem. 85, 5288–5296 (2013).

15. Hu, Q. et al. The Orbitrap: a new mass spectrometer. J. Mass Spectrom. 40, 430–443 (2005).

16. Kelstrup, C. D., Young, C., Lavallee, R., Nielsen, M. L. & Olsen, J. V. Optimized fast and sensitive acquisition methods for shotgun proteomics on a quadrupole orbitrap mass spectrometer. J. Proteome Res. 11, 3487–3497 (2012).

17. Michalski, A. et al. Mass spectrometry-based proteomics using Q exactive, a high-performance benchtop quadrupole orbitrap mass spectrometer. Mol. Cell. Proteomics 10, M111.011015 (2011).

18. Kelstrup, C. D. et al. Rapid and deep proteomes by faster sequencing on a benchtop quadrupole ultra-high-field orbitrap mass spectrometer. J. Proteome Res. 13, 6187–6195 (2014).

19. Trujillo, E. A., Hebert, A. S., Brademan, D. R. & Coon, J. J. Maximizing Tandem Mass Spectrometry Acquisition Rates for Shotgun Proteomics. Anal. Chem. 91, 12625–12629 (2019).

20. Grinfeld, D., Aizikov, K., Kreutzmann, A., Damoc, E. & Makarov, A. Phase-constrained spectrum deconvolution for fourier transform mass spectrometry. Anal. Chem. 89, 1202– 1211 (2017).

21. Lange, O., Damoc, E., Wieghaus, A. & Makarov, A. Enhanced Fourier transform for Orbitrap mass spectrometry. Int. J. Mass Spectrom. 369, 16–22 (2014).

22. Steigerwald, S. et al. Full Mass Range ΦSDM Orbitrap Mass Spectrometry for DIA Proteome Analysis. Mol. Cell. Proteomics 23, 100713 (2024).

23. Kelstrup, C. D. et al. Limits for Resolving Isobaric Tandem Mass Tag Reporter Ions Using Phase-Constrained Spectrum Deconvolution. J. Proteome Res. 17, 4008–4016 (2018).

24. Bekker-Jensen, D. B. et al. An Optimized Shotgun Strategy for the Rapid Generation of Comprehensive Human Proteomes. Cell Syst. 4, 587-599.e4 (2017).

25. Kelstrup, C. D. et al. Performance Evaluation of the Q Exactive HF-X for Shotgun Proteomics. J. Proteome Res. 17, 727–738 (2018).

26. Fröhlich, K. et al. Benchmarking of analysis strategies for data-independent acquisition proteomics using a large-scale dataset comprising inter-patient heterogeneity. Nat. Commun. 13, (2022).

27. Nagaraj, N. et al. System-wide perturbation analysis with nearly complete coverage of the yeast proteome by single-shot ultra HPLC runs on a bench top orbitrap. Mol. Cell. Proteomics 11, (2012).

28. Makarov, A. & Denisov, E. Dynamics of Ions of Intact Proteins in the Orbitrap Mass Analyzer. J. Am. Soc. Mass Spectrom. 20, 1486–1495 (2009).

29. Kafader, J. O. et al. Measurement of Individual Ions Sharply Increases the Resolution of Orbitrap Mass Spectra of Proteins. Anal. Chem. 91, 2776–2783 (2019).

30. Batth, T. S. et al. Protein aggregation capture on microparticles enables multipurpose proteomics sample preparation. Mol. Cell. Proteomics 18, 1027–1035 (2019).

31. Müller, T. et al. Automated sample preparation with SP 3 for low-input clinical proteomics. Mol. Syst. Biol. 16, 1–19 (2020).

32. Cox, J. & Mann, M. MaxQuant enables high peptide identification rates, individualized p.p.b.-range mass accuracies and proteome-wide protein quantification. Nat. Biotechnol. 26, 1367–1372 (2008).

33. Wickham, H. et al. Welcome to the Tidyverse. J. Open Source Softw. 4, 1686 (2019).

34. Villanueva, R. A. M. & Chen, Z. J. ggplot2: Elegant Graphics for Data Analysis (2nd ed.). Meas. Interdiscip. Res. Perspect. 17, 160–167 (2019).

35. Kockmann, T. & Panse, C. The rawrr R Package: Direct Access to Orbitrap Data and beyond. J. Proteome Res. 20, 2028–2034 (2021).

